# GDTR: Layer-wise Settling Depth Reveals Biological Grammar in Genomic Foundation Models

**DOI:** 10.64898/2026.07.14.738370

**Authors:** Yoonjin Cho, Jiheon Kang, Subin Park, Sangwoo Kim

**Affiliations:** Yonsei University College of Medicine, Seoul, Korea; Department of Electrical and Electronic Engineering, Yonsei University, Seoul, Korea; Department of Computer Science, Yonsei University, Seoul, Korea; Department of Biomedical Systems Informatics, Yonsei University College of Medicine, Seoul, Korea

## Abstract

Genomic foundation models capture sequence regularities, yet existing interpretability tools rarely ask *where* in the layer stack a biological grammar becomes stable. We introduce GDTR (*Genomic Deep-Thinking Ratio*), a training-free residual-stream lens that assigns each nucleotide token a *settling depth c*(*t*): the first layer at which its representation stabilises against the post-final-norm reference. On Evo 2 7B, splice donor and acceptor sites settle approximately two layers earlier than intronic contexts, enhancer-like cCREs show a smaller but measurable shift, and a chr22 calibration transfers to held-out chr17. Perturbing canonical splice donors shows that the signal is bidirectional: disrupting the central GT motif deepens settling, whereas shuffling the flanking grammar makes the preserved motif settle earlier. Differential GDTR further reveals consequence-associated peak-disruption depths across ClinVar variants, with synonymous substitutions peaking deepest but with broad class overlap. GDTR therefore provides a layer-wise interpretability axis for genomic foundation models, complementary to existing prediction and variant-scoring tools.

## 1. Introduction

Genomic foundation models (Nguyen et al., 2026; 2023; Dalla-Torre et al., 2024; Zhou et al., 2024; Schiff et al., 2024) are increasingly used across functional-genomics tasks, but their internal layer-wise dynamics remain difficult to interpret. Existing tools ask *where* models attend (Abnar & Zuidema, 2020; Avsec et al., 2021), *what* features they encode (Bricken et al., 2023; Cunningham et al., 2023), or *how* variants shift final representations (Cheng et al., 2023; Jaganathan et al., 2019). Even recent sparse-autoencoder analyses of Evo 2 (Nguyen et al., 2026; Deng et al., 2025) recover interpretable genomic features from selected layers, answering *what* the model represents. A complementary question remains largely unmeasured: *when, along the layer stack, does a biological sequence element become stable?* On Evo 2, a splice donor’s representation stabilises about two layers earlier than a nearby intronic position, even though both end at the same final layer.

We introduce GDTR (*Genomic Deep-Thinking Ratio*), a training-free residual-stream lens (a per-layer readout of the model’s hidden state) for this question. GDTR assigns each nucleotide token a *settling depth c*(*t*): the first layer at which its representation stabilises against the model’s output-ready reference. We use *biological grammar* to mean the context-dependent integration of a local motif with its flanking sequence, as in splice donors where the central GT dinucleotide is functional only in context. GDTR adapts the Deep-Thinking Ratio idea (Chen et al., 2026) to genomic causal LMs with a cosine residual-stream lens, a running-minimum envelope that smooths non-monotone layer-by-layer trajectories, and an Evo 2-specific post-final-norm reference. Fig. 1 summarises the readout; details are in §2.

**Figure 1.**
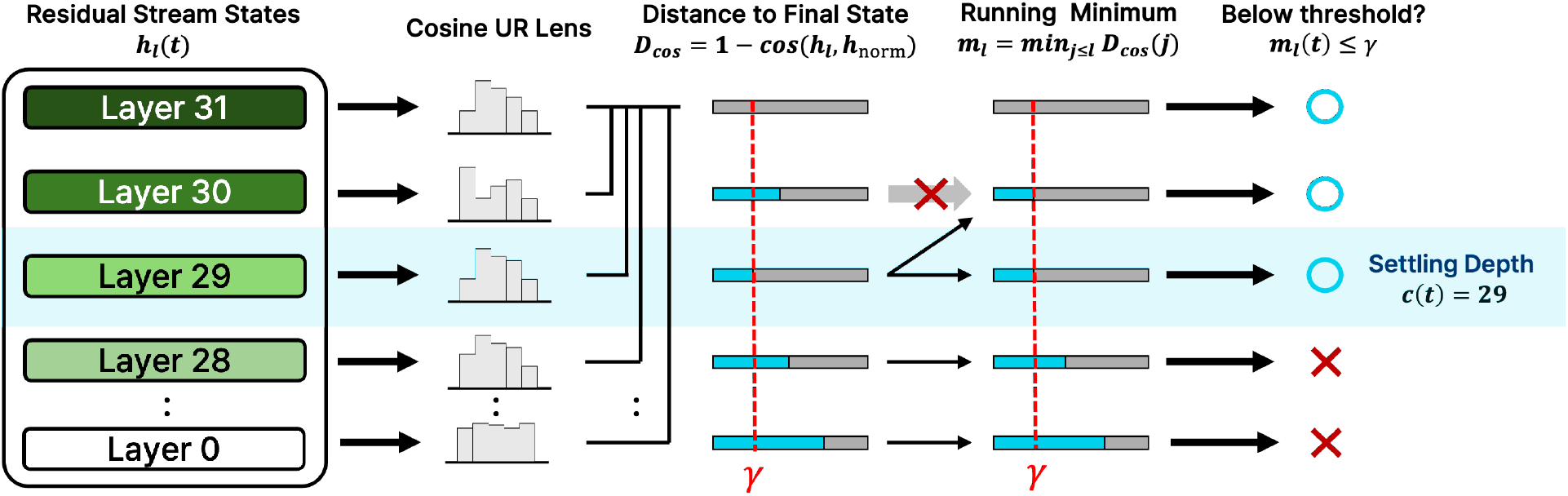
GDTR measures the layer at which a token’s representation settles into the model’s output-ready frame. Each Evo 2 layer emits a residual-stream tap *h*_ℓ_(*t*); the *cosine lens* compares it to the post-final-norm reference *h*_norm_ via *D*_cos_(*ℓ, t*) = 1*−*cos(*h*_ℓ_(*t*), *h*_norm_(*t*)) (*distance to final state*). The *running-minimum envelope m*_ℓ_ =min_j*≤*ℓ_ *D*_cos_(*j*) enforces a monotone non-increasing trace, and the settling depth *c*(*t*) is the first (shallowest) layer at which *m*_ℓ_ ≤ *γ* (Eq. 2). In the illustrative trace shown, *c*(*t*) = 29: a raw distance at a later layer rises back above *γ*, but the envelope carries the smaller value at layer 29 forward, so every layer at or beyond layer 29 satisfies *m*_ℓ_ ≤ *γ* while shallower layers (*ℓ* ≤ 28) do not. Lower *c*(*t*) means earlier stabilisation. The layer numbers in this schematic are pedagogical; Evo 2’s actual layer-30 saturation is discussed in §2 and App. A.1.

Our central claim is that settling depth is biologically structured. On Evo 2 7B, a single chr22 calibration of *c*(*t*) transfers to held-out chr17; splice sites and enhancer-like cCREs settle earlier than surrounding contexts; and matched motif and flank perturbations on the same donor loci move settling depth in opposite directions, separating motif detection from flanking-context integration. Subtracting the reference-allele cosine trajectory from the alternate-allele one, the same lens shows consequence-associated peak-disruption depths across ClinVar variant classes. GDTR therefore provides a layer-resolved interpretability axis for genomic foundation models, complementary to final prediction scores and variant scorers.

## 2. The GDTR Framework

For a nucleotide sequence of length *T* processed by a causal language model with *L* layers, we extract the residual-stream activation *h*_ℓ_(*t*) at every layer *ℓ* ∈{1, …, *L* }and token position *t*. We replace the JSD lens of the parent DTR (Chen et al., 2026) with a cosine residual-stream lens that operates directly in hidden-state geometry,

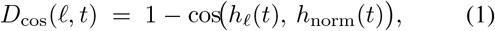

where *h*_norm_(*t*) is the post-final-RMSNorm state that the model’s unembedding head reads (layers indexed 0, …, *L* −1 throughout the Evo 2 analysis). Evo 2 tokenises at single-nucleotide (1-mer) resolution, so its next-token mass concentrates on an approximately 4-symbol alphabet, and the parent DTR’s logit-lens JSD collapses to a near-binary trace with little discriminative resolution. The cosine distance in the 4096-dimensional residual stream descends smoothly enough to threshold into *c*(*t*) and avoids the unembedding projection, which is unreliable through Evo 2’s rotations and pre-final saturation (App. A.1). We retain JSD only as a cross-lens check for variant discrimination (App. C), where aggregation over positions restores its signal. We define the *settling depth* via a running-minimum envelope,

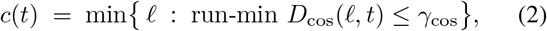

where run-min *D*_cos_(*ℓ, t*) = min_k*≤*ℓ_ *D*_cos_(*k, t*). Note that this is equivalent to the first-passage time min {*ℓ* : *D*_cos_(*ℓ, t*) ≤ *γ*_cos_ }; the running-minimum is written explicitly to reflect the monotone envelope used for visualization in Fig. 1.

Lower *c*(*t*) means earlier stabilisation; higher *c*(*t*) means that the token remains unresolved until later layers. NLP transformers exhibit near-monotonic logit-lens convergence (nostalgebraist, 2020; Belrose et al., 2023; Pal et al., 2023); genomic CLMs do not (Hyena alignment rotations, Evo 2 pre-final saturation), so the running-min reduces this non-monotonicity to a monotone-non-increasing trace. The threshold *γ*_cos_ is set by *regional q*_70_ *calibration*: the 70th percentile of the running-min at the penultimate layer over the analysis region. The operating point sits within a wide, flat plateau identified by a 5 × 5 grid search over (*γ*_cos_, *ρ*) (App. A.2).

### Settling depth is two-sided

A lower *c*(*t*) does not by itself imply a stronger biological signal. A token can settle early because (a) its representation is constrained by a biological grammar that the model commits to early, *or* (b) its surrounding context is simpler, so the running-min envelope reaches *γ*_cos_ with little integration. We exploit this two-sidedness directly: the perturbation experiments in §3.2 are designed to push *c*(*t*) in opposite directions, separately stressing motif detection and flanking-grammar integration. Throughout the paper we therefore interpret depth signatures as *bidirectional*. Both directions of movement can carry information about motif detection and context integration, but they should not be read as a monotone scale of biological signal strength. As a concrete instance of arm (b), non-canonical splice donors settle *earlier* than the dominant GT-AG class (App. D.2), reflecting tighter flanking constraint rather than a stronger splice signal.

### Representational saturation before the final layer

The reference *h*_norm_ is fixed by the model as the post-final-RMSNorm state that the unembedding reads, independently of any tap we select. Measured against it, Evo 2 reaches the post-norm direction before the last attention layer executes: on chr22 control sequences, max_t_|*h*_30_(*t*) − *h*_31_(*t*) |= 0 exactly (layer 31 is a residual passthrough), while cos(*h*_29_, *h*_norm_) = −0.013 versus cos(*h*_30_, *h*_norm_) = cos(*h*_31_, *h*_norm_) =+0.6855. Layer 30 is thus the step that rotates the representation into the output-ready frame, leaving layer 29 as the deepest tap not yet aligned with it. Since the running-min envelope (Eq. 2) records how early a token reaches *h*_norm_, the discriminating variation in settling depth lies in these pre-rotation layers. We report *L*^⋆^ = 29 as that empirical boundary rather than as a claim about the model’s architectural design intent; full evidence is in App. A.1.

### Tuned-lens consistency check against frame mismatch

Frame mismatch is the worry that interior layers sit in a different linear basis from *h*_norm_, so a raw cosine distance would track basis rotation rather than genuine convergence. A per-layer tuned-lens affine fit (Belrose et al., 2023) reaches ≥ 98% MSE recovery on 30*/*32 Evo 2 layers (canonical *L*^⋆^ = 29: 0.9996; App. A.4), so the interior taps are one affine transform away from the output frame. Gross frame mismatch is therefore unlikely to drive the GDTR signal. No affine weights are used at inference, so the framework remains training-free.

## 3. Results

### Statistical conventions

The test in each subsection is fixed by the structure of its data rather than chosen for convenience: §3.1 uses the Mann–Whitney *U* test (independent two-sample comparisons at large *n*), §3.2 uses the paired Wilcoxon signed-rank test (the same donor loci under matched perturbations), and §3.3 uses a Kruskal–Wallis omnibus with Bonferroni-corrected Dunn post-hoc tests (three or more unmatched groups of discrete, tied argmax layers). We report Cohen’s *d* wherever comparisons are pairwise, *ε*^2^ for the Kruskal–Wallis omnibus, and the rank-biserial *r* for Dunn contrasts, so that effect magnitudes remain comparable across subsections.

### 3.1. Splice and Enhancer Annotations Settle Early

We first test whether *c*(*t*) produces a transferable readout at the genome scale. Chromosome 22 is the calibration set (*γ*_cos_ = 0.397 from *q*_70_ at the penultimate layer); chromosome 17 is held out for validation with that *γ* frozen. The chr17 *q*_70_ recomputed independently equals 0.394 (|Δ| = 0.0023); the chr17 splice-donor effect reaches 94.6% of the chr22 magnitude (Cohen’s *d* = −0.349 vs intron), and the relative ordering of the seven canonical contexts is preserved (Spearman *ρ* = 0.93). The calibration transfers without re-tuning, and we use this single *γ* for all subsequent context- and variant-level comparisons. All context-level summaries below report the per-position *c*(*t*) pooled over the analysis windows, summarised by Cohen’s *d* versus intron, which normalises by the pooled within-context standard deviation and so absorbs within-region variability by construction.

### Context-level ordering

On canonical genomic contexts (Fig. 2a), splice donor and acceptor sites settle approximately 2 layers earlier than the intronic baseline, and the remaining contexts cluster near or above it. Given the large number of pooled positions (≈10^7^), we treat effect size rather than nominal significance as the primary evidence and Mann–Whitney *p*-values as a secondary check. Among the regulatory annotations (Fig. 2b), all Cohen’s *d* values are computed against the same pooled chr17+chr22 intron baseline 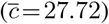 used in panel (a), so cross-panel magnitudes are directly comparable. ENCODE cCRE-ELS enhancer-like elements (Moore et al., 2020) are the clearest non-splice signal at Cohen’s *d* = −0.118 on this baseline (vs. splice-donor *d* = −0.354): biologically modest but statistically robust, and the effect transfers from chr22 to chr17. SNP-level annotations (GTEx Whole-Blood chr22 eQTLs (The GTEx Consortium, 2020): *n* = 32,456, *d* =+0.064; GWAS Catalog chr22 SNPs (Sollis et al., 2023): *n* = 6,900, *d* =+0.021) yield negligible effect sizes that we do not interpret further. The 5^*′*^ UTR shift is reported separately because it showed anomalous entropy coupling on the 120-window control panel (*ρ* = +0.41), although this coupling attenuates substantially at full-chr22 scale (*ρ* = +0.054, with |*ρ*| ≤0.224 across all seven canonical contexts; App. E). We keep 5^*′*^ UTR on a separate axis, a conservative carryover from the smaller panel.

**Figure 2.**
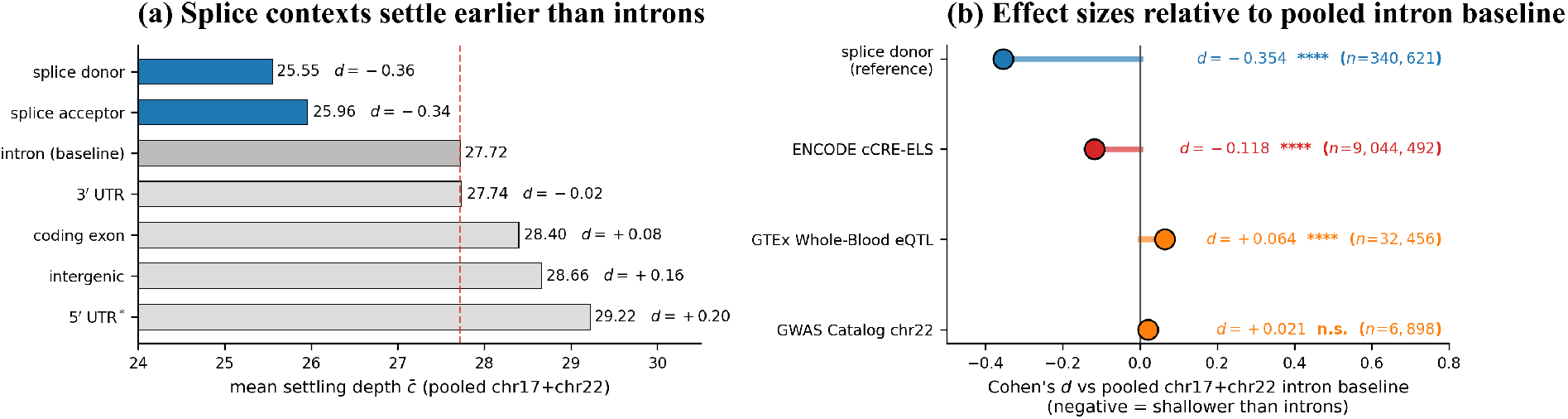
Splice and enhancer annotations shift toward shallower settling. Both panels report Cohen’s *d* against the pooled chr17+chr22 intron baseline (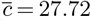, red dashed). **(a)** Per-context mean settling depth 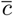 on canonical genomic contexts (frozen *γ*_cos_ = 0.397). Splice donor and acceptor (blue) sit approximately 2 layers below the intron baseline. The asterisk on “5^*′*^ UTR^*\**^” marks an entropy-coupled context (§4) reported separately from the entropy-controlled hierarchy. Right of each bar: Cohen’s *d* versus the intron baseline (Mann–Whitney significance: *,**,***,**** at 0.05, 0.01, 10^*−*3^, 10^*−*4^). **(b)** Cohen’s *d* for regulatory annotations: splice donor (reference; *−*0.354), ENCODE cCRE-ELS (*−*0.118), GTEx Whole-Blood eQTL (+0.064), GWAS Catalog (+0.021). cCRE-ELS yields a measurable enhancer signal; SNP-level eQTL and GWAS shifts are negligible on the effect-size scale.

### Entropy does not explain the splice signal

A natural concern is that *c*(*t*) merely tracks the per-position next-token Shannon entropy *H*_t_ of the model. On a control panel of 120 random chr22 windows (719,000 analysed positions) we compute *H*_t_ from the post-norm logits and find an overall Spearman *ρ*(*c, H*_t_) = −0.079 (per-context |*ρ*| ≤0.16 except 5^*′*^ UTR; §4; full per-context numbers in Tab. A3, App. A.3). On the same chr22 panel the splice-donor-versus-intron Cohen’s *d* is − 0.452; after regressing *c* on *H*_t_ and re-measuring on the residual it strengthens to −0.583. Next-token entropy therefore does not explain the splice depth ordering.

### 3.2. Motif Edits Separate Detection from Flanking Context

We probe *c* (*t*) with two complementary perturbations on the same 1,000 canonical GT-AG donors (chr22, ±10 bp pad-averaged 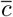). Real donors settle at 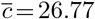 (Table 1). Replacing the central GT with AA while preserving the ±100 bp flank deepens 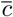 by 0.46 layers (paired Wilcoxon *p* = 2.3 × 10^*−*32^, |*d*| = 0.086): a small but reliable signal that the model uses the canonical dinucleotide for early commitment. The GT →AA edit also changes composition; a composition-preserving variant (e.g. GT→ TG) would isolate motif identity from composition. Dinucleotide-shuffling the ±100 bp flank while preserving the central GT produces the opposite shift, lifting *c* to a shallower value by 3.18 layers (paired *p* = 4.1 ×10^*−*59^, |*d*| = 0.515): the isolated GT becomes easier to stabilise, whereas the real donor context requires deeper flanking-grammar integration. The two perturbations move *c* in opposite directions on the same donor loci, separating motif detection from context integration along the same axis. We cannot rule out from depth alone an alternative reading under which the model *disengages* from uninformative noise on shuffled flanks and commits early to the lone GT (“early simplification” rather than reduced integration); the within-splice motif breakdown in App. D.2 carries the same cautionary logic.

**Table 1.**
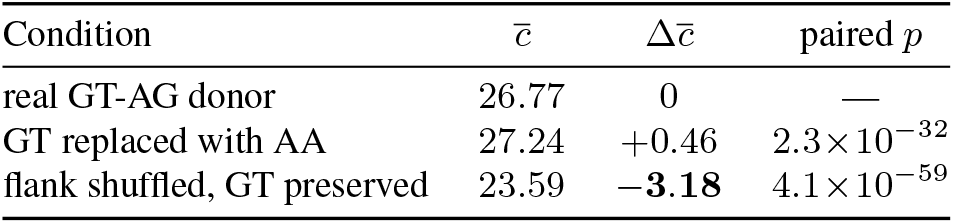
Motif-edit and flank-shuffle controls on 1,000 canonical GT-AG donors (chr22, ±10 bp pad-averaged 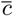). Paired Cohen’s |*d*| is 0.086 for the GT motif edit and 0.515 for the flank shuffle.

| Condition | $\bar{c}$ | $\Delta\bar{c}$ | paired $p$ |
| --- | --- | --- | --- |
| real GT-AG donor | 26.77 | 0 | — |
| GT replaced with AA | 27.24 | +0.46 | $2.3 \times 10^{-32}$ |
| flank shuffled, GT preserved | 23.59 | −3.18 | $4.1 \times 10^{-59}$ |

### 3.3. Variant-Disruption Depth Correlates with Molecular Consequence

For variants we read the same cosine trajectory differentially as 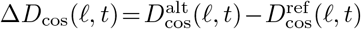 and record the layer at which |Δ*D*_cos_| peaks. This converts the GDTR lens from an absolute settling readout into a sensitivity readout. Across 4,023 ClinVar P/LP variants in 15 cancer-associated genes, molecular-consequence classes show different peak-disruption distributions, with class medians ordered *intron < frameshift < nonsense < missense* ≈ *canonical splice < synonymous* (Fig. 3). The Kruskal–Wallis omnibus is significant (*p* = 3.0 × 10^*−*10^, *H* = 53.2) (Kruskal & Wallis, 1952), but the effect is small (*ε*^2^ = 0.013) and the class distributions overlap broadly; only the nonsense-versus-missense contrast is individually resolved after Bonferroni-corrected Dunn testing (*p*_adj_ *<* 10^*−*3^) (Dunn, 1964). Pathogenic synonymous variants often act through splicing, so the synonymous tail may carry the canonical-splice signal rather than a separate synonymous one. We read this as a population-level depth shift rather than class separation or a pathogenicity score. Full Δ*D*_cos_ trajectory AUROC analyses, cohort-construction details, and rank-biserial effect sizes are in App. C.

**Figure 3.**
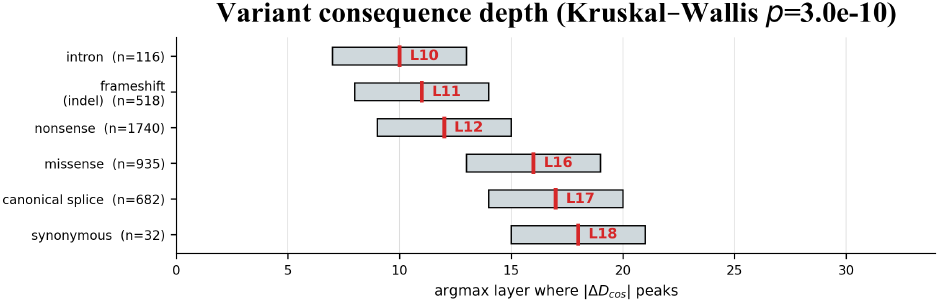
Variant consequences peak at different disruption layers. Argmax layer of Δ*D*_cos_ across 4,023 ClinVar P/LP variants in 15 cancer-associated genes, grouped by molecular consequence; class-median layer in red. Kruskal–Wallis 6-way *p* = 3.0*×* 10^*−*10^; the largest adjacent jump is between nonsense and missense (*p*_adj_ *<* 10^*−*3^). The broad overlap reflects a population-level depth shift rather than class separation.

### 3.4. Robustness and Tokenisation Limits

Applying the same chr22 protocol to Evo 2 7B, HyenaDNA-large, NT-v2 500M, and DNABERT-2 gives a bounded robustness result (Table 2; details in App. B). All four baselines use official HuggingFace checkpoints with frozen weights and the same regional *q*_70_ calibration protocol.

**Table 2.** Cross-model summary. Full per-window numbers and MLM tokenisation notes are in App. B.

| Family (model) | Per-pos. readout | Donor < intron |
| --- | --- | --- |
| Causal LM (Evo 2, HyenaDNA-large) | yes | yes |
| MLM (NT-v2, DNABERT-2) | no (per-window) | resolution-limited |

Donor settles earlier than intron in both per-base causal LMs, and their per-window settling depths are moderately concordant across models (Spearman *ρ* = +0.516). The MLMs operate at *k*-mer or BPE token granularity and cannot resolve single-base splice junctions. The cross-model result supports GDTR within per-position causal LMs while identifying tokenisation granularity as the current architectural limit.

## 4. Discussion and Limitations

Settling depth is bidirectional, and this has a cost as well as a use. The use is that it explains within-class orderings that run against naive canonicity priors (Apps. D.2, F) and warns against reading *c*(*t*) as a scale of biological signal strength. The cost is that a single *c*(*t*) does not interpret itself: a shallow settling can mean early motif commitment, a highly determinate context, or disengagement from uninformative input. The two matched perturbations of §3.2 move *c* in opposite directions: disrupting the central GT deepens *c*, whereas shuffling the flanking grammar makes the preserved GT settle earlier. The context arm moves *c* about six times as far as the motif arm (|*d*| = 0.515 vs 0.086), so on those loci *c*(*t*) behaves mainly as a contextual-determinacy readout; whether the same balance holds at other functional contexts is open.

The remaining limitations group into four kinds. The lens is direction-only and discards residual magnitude (App. G.1), and the absolute readout saturates past the rotation layer, so settling and variant scoring are not read from the same layers (§3.3, App. C). The motif and flank edits of §3.2 are off-manifold and cannot, on their own, separate reduced integration from early simplification; the within-splice ordering in App. D.2 is the on-manifold version of the same test. Composition and *k*-mer rarity are only partly controlled (App. G.2): the splice effect survives entropy and GC matching, but promoter and flank-shuffle contrasts remain composition-entangled (App. F). Scope is narrow: a single calibration chromosome transferred to one held-out chromosome, replication on the only other per-base model available (HyenaDNA; Caduceus (Schiff et al., 2024) is the obvious next per-base architecture to test), and a variant cohort of 15 cancer-associated genes.

The variant-depth ordering hints at an analogy with hierarchical encoding in protein language models (Banerjee et al., 2025), but encoding depth, settling depth, and sensitivity-peak depth are distinct quantities, so we treat this as a hypothesis rather than a result. A concrete next step is to test it against public Evo 2 sparse autoencoders (Deng et al., 2025): do early-settling positions align with the activations of specific splice or exon SAE features? The rotation boundary itself is a portable methodological result. A residual-stream lens taken from NLP needs both a per-model reference and an explicit choice between absolute and differential readouts on genomic models (§2, App. C), alongside causal or sparse-feature interventions (Cunningham et al., 2023; Meng et al., 2022) that connect depth signatures to model circuits.

## 5. Conclusion

GDTR provides a training-free, layer-resolved probe of where genomic foundation models stabilise token representations relative to their output-ready residual frame. Across Evo 2 analyses, settling depth is chromosome-transferable, shifts earlier at splice sites and enhancer-like cCREs, and responds in opposite directions to motif edits and flank shuffles, indicating that depth reflects both motif detection and contextual integration. Differential GDTR further shows that variant-induced residual disruptions peak at consequence-associated layers, suggesting a complementary axis for interpreting variant effects rather than a standalone pathogenicity score. Future work should extend the analysis to whole-genome matched controls, additional per-base architectures, and causal or sparse-feature interventions (Cunningham et al., 2023; Meng et al., 2022) that connect depth signatures to concrete model circuits.

## Appendix

*Supplementary material for “*GDTR: *Layer-wise Settling Depth Reveals Biological Grammar in Genomic Foundation Models”*

### A. Method Details

#### A.1. Addressing the Structural Characteristics of Evo 2: A Non-Functional Final Layer

We observe that the final attention layer of Evo 2 7B is structurally non-functional with respect to the residual stream. Direct measurement on chr22 control sequences (100 windows of 6 kb each) yields the values in Table A1. Two observations are relevant: (i) max_t_ |*h*_30_(*t*) − *h*_31_(*t*) |= 0 exactly, so layer 31 acts as a residual passthrough; (ii) the cosine alignment with the post-final-norm reference *h*_norm_ shifts from cos(*h*_29_, *h*_norm_) = −0.013 (near-orthogonal) to cos(*h*_30_, *h*_norm_) = cos(*h*_31_, *h*_norm_) = +0.6855, indicating that layer 30 performs the rotation into the post-norm direction. Consequently, the deepest tap not yet aligned with the output-ready frame is *L*^⋆^ = 29. The standard NLP convention of tapping the final layer therefore does not apply to Evo 2; we instead use the post-final-RMSNorm state *h*_norm_ as the convergence reference and report *L*^⋆^ = 29 as the empirical boundary of the pre-rotation taps.

**Table A1.**
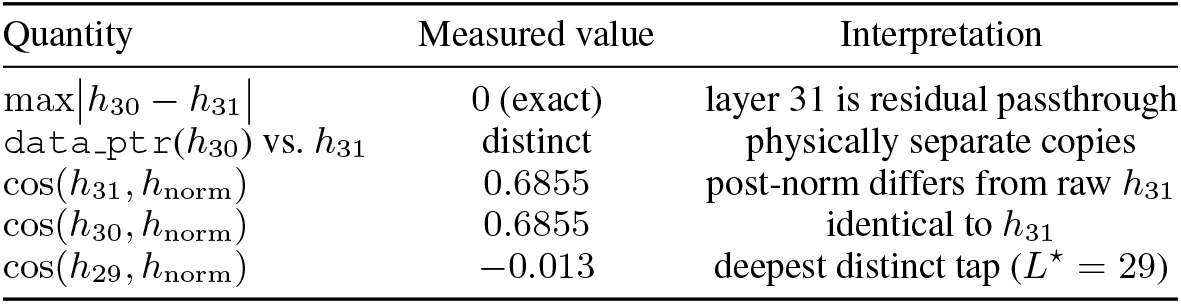
Evidence that the final attention layer of Evo 2 is structurally non-functional with respect to the residual stream. Layer 31 is bit-identical to layer 30 in value but stored as a distinct tensor; both align cosine-wise with the post-final-RMSNorm reference, identifying layer 29 as the deepest tap not yet aligned with the output frame.

| Quantity | Measured value | Interpretation |
| --- | --- | --- |
| $\max \vert h_{30} - h_{31}\vert $ | 0 (exact) | layer 31 is residual passthrough |
| $\text{data\_ptr}(h_{30})$ vs. $h_{31}$ | distinct | physically separate copies |
| $\cos(h_{31}, h_{\text{norm}})$ | 0.6855 | post-norm differs from raw $h_{31}$ |
| $\cos(h_{30}, h_{\text{norm}})$ | 0.6855 | identical to $h_{31}$ |
| $\cos(h_{29}, h_{\text{norm}})$ | -0.013 | deepest distinct tap ($L^* = 29$) |

##### Why the rotation matters as a measurement boundary

This boundary has two consequences for the framework. First, the standard NLP recipe of reading the final layer would compare against a passthrough and an already-saturated state, which collapses the signal, so the post-final-norm reference is needed rather than cosmetic. Second, it places the settling readout in the pre-rotation taps (*ℓ* ≤ 29). Once layer 30 aligns the stream with *h*_norm_, the absolute cosine distance saturates and no longer separates tokens. That is a property of the absolute readout, not evidence that later layers are inert: read *differentially*, layer 30 carries the sharpest variant signal (§3.3, App. C). The rotation is therefore a boundary in measurement geometry, with level readouts useful before it and sensitivity readouts after it. It is also the main reason a residual-stream lens taken from NLP needs a per-model reference on genomic models.

#### A.2. Hyperparameter Sensitivity

On chr22 the regional *q*_70_ calibration yields *γ*_cos_ = 0.397. A 5×5 grid search over (*γ*_cos_, *ρ*) (Table A2) produces a wide, flat plateau around the operating point (*γ*_cos_ = 0.40, *ρ* = 0.85) with ±0.10× ±0.05 variation. Within this plateau the standard deviation across the 25 cells is 0.06, indicating low sensitivity to small perturbations in either hyperparameter. This robustness was first identified in Phase 0 (HyenaDNA-medium-160k), where the analogous *q*_70_ calibration produced a best operating point of (*γ*_cos_ = 0.50, *ρ* = 0.85) with Cohen’s *d* = −1.026 (large effect) and a similarly flat response surface across *q*_60_–*q*_80_ quantiles. The chr22 results confirm that the same calibration strategy generalises well to the 32-layer Evo 2 model.

**Table A2.** Effect-size grid search over (*γ*_cos_, *ρ*). Values are reported on the analysis scale used for hyperparameter calibration. The locked operating point (0.40, 0.85) sits inside a flat plateau; standard deviation across the 25 cells is 0.06.

| $\gamma_{\text{cos}} \backslash \rho$ | $\rho = 0.70$ | $\rho = 0.75$ | $\rho = 0.80$ | $\rho = 0.85$ | $\rho = 0.90$ |
| --- | --- | --- | --- | --- | --- |
| 0.30 | 5.04 | 5.18 | 5.21 | 5.20 | 5.15 |
| 0.35 | 5.10 | 5.21 | 5.24 | 5.23 | 5.18 |
| 0.40 | 5.16 | 5.25 | **5.28** | **5.26** | 5.21 |
| 0.45 | 5.13 | 5.22 | 5.25 | 5.24 | 5.19 |
| 0.50 | 5.08 | 5.17 | 5.20 | 5.19 | 5.14 |

#### A.3. Entropy Decoupling per Context

Table A3 reports per-context Spearman *ρ*(*c, H*_t_) on a 120-window chr22 control panel (719,000 analysed positions; the 1,000 shortfall versus 120 × 6,000 reflects truncated logits at window edges where the causal context is incomplete), where *H*_t_ is the per-position next-token Shannon entropy from the post-norm logits. The overall correlation is small and slightly negative (*ρ* =−0.079); per-context correlations stay |*ρ*|≤ 0.16 for every context except 5^*′*^ UTR (*ρ* =+0.41). The largest non-UTR couplings are splice acceptor at |*ρ*| = 0.152 and intron at |*ρ*| = 0.108 (Tab. A3). After regressing *c* on *H*_t_ on the same chr22 panel, the splice-donor-versus-intron Cohen’s *d strengthens* from −0.452 to −0.583 on the residual, confirming that next-token entropy does not explain the splice depth ordering.

**Table A3.** Per-context Spearman correlation between settling depth *c*(*t*) and next-token entropy *H*_t_ on the chr22 control panel (120 windows, 719,000 analysed positions; the 1,000 shortfall versus 120*×*6,000 reflects truncated logits at window edges). The next-largest non-UTR couplings are splice acceptor (|*ρ*| = 0.152) and intron (|*ρ*| = 0.108); 5^*′*^ UTR is the only entropy-coupled context.

| Context | $n$ | $\bar{c}$ | $\bar{H}_t$ | $\rho(c, H_t)$ |
| --- | --- | --- | --- | --- |
| intergenic | 243,956 | 28.94 | 0.795 | $-0.024$ |
| intron | 425,131 | 28.03 | 1.030 | $-0.108$ |
| coding exon | 36,427 | 28.61 | 0.744 | $-0.080$ |
| 3’ UTR | 8,238 | 27.61 | 1.073 | $-0.028$ |
| splice donor | 1,981 | 25.04 | 0.887 | $-0.083$ |
| splice acceptor | 1,920 | 27.33 | 0.764 | $-0.152$ |
| **5’ UTR** | 2,347 | 31.43 | 1.121 | **$+0.414$** |
| overall | 719,000 | — | — | $-0.079$ |

#### A.4. Tuned-Lens Recovery Across All 32 Layers

Each layer is fitted with a single 4096 × 4096 affine *A*_ℓ_ using MSE between *A*_ℓ_*h*_ℓ_ and *h*_norm_, optimised with Adam (*β*_1_ = 0.9, *β*_2_ = 0.999, weight decay = 0) at learning rate 10^*−*3^ for 15 epochs over 100 calibration sequences with batch size 1. The affine weights are used only for this verification and are discarded at inference, so no hyperparameter governs the production settling-depth pipeline. The post-norm tap is the prediction target. Recovery scores at five representative layers are reported in Table A4; across the full 32-layer distribution, 30 reach ≥ 98%, with the two layers below this threshold not among the five shown (so the table’s “worst of these five” label refers to the displayed subset, not the overall minimum). The ≥ 98% threshold is descriptive of the empirical recovery distribution rather than a pre-registered cut-off: it summarises where the bulk of the per-layer recovery scores sit, not a criterion that the framework is required to clear at inference (the framework is training-free and uses no affine weights when computing *c*(*t*)).

**Table A4.** Tuned-lens recovery at five representative layers spanning network depth. Block-type abbreviations follow Evo 2’s Striped-Hyena 2 architecture: hcl (Hyena-Convolution-Long), hcm (Hyena-Convolution-Medium), hcs (Hyena-Convolution-Short).

| Layer / block type | Initial MSE | Final MSE (15 epochs) | Recovery |
| --- | --- | --- | --- |
| $\ell = 2$ (hcl) | 1,259 | 5.6 | 0.9956 |
| $\ell = 12$ (hcm) | 742 | 13.6 | 0.9816 (worst of these five) |
| $\ell = 22$ (hcm) | 418 | 0.41 | 0.9990 |
| $\ell = 28$ (hcs) | 822 | 0.34 | 0.9996 (best below tap) |
| $\ell = 29$ (canonical tap, hcm) | 510 | 0.20 | 0.9996 |

### B. Cross-Architecture Replication: Scope, Granularity, Two-Tier Structure

This appendix expands the bounded robustness result of §3.4. We compare four publicly released genomic foundation models that span the per-bp causal-LM and the tokenised MLM design space: Evo 2 7B, HyenaDNA-large-1m, NT-v2 500M, and DNABERT-2 117M. Architectural specifications, the per-window token budget they impose on the same chr22 sequences, end-to-end runtime, and the per-model *q*_70_-calibrated *γ*_cos_ are summarised in Table A5. All four models use the identical

**Table A5.** Architectural specifications, tokenisation, per-window context, runtime, and per-model *q*_70_-calibrated *γ*_cos_ for the four genomic foundation models compared in §3.4.

| Model | Architecture | Layers | Hidden | Tokens / 6 kb window | Wall-clock | $\gamma_{q_{70}}$ |
| --- | --- | --- | --- | --- | --- | --- |
| Evo 2 7B | Hybrid Transformer + StripedHyena 2 | 32 | 4096 | 6,000 (1 bp) | reused | 0.397 |
| HyenaDNA-large-1m | Pure Hyena | 8 | 256 | 6,001 (1 bp + BOS) | $\approx 4$ min | 0.358 |
| NT-v2 500M | Transformer MLM ($k$-mer) | 29 | 1024 | 671 ($k = 6$, $\approx 4$ kb) | $\approx 7.5$ min | 0.533 |
| DNABERT-2 117M | Transformer MLM (BPE) | 12 | 768 | $\approx 600$ (BPE, $\approx 3$ kb) | $\approx 3$ min | 0.677 |

#### Per-context replication

Table A6 reports the per-context mean settling depth for each model. The two per-bp causal LMs (Evo 2, HyenaDNA-large) replicate donor settles earlier than intron, and acceptor earlier than intron, at depth-normalised scale (approximately 3 layers below intron in 32-layer Evo 2; approximately 0.34 layers below intron in 8-layer HyenaDNA). The two MLMs tokenise at *k*-mer/BPE granularity: their per-window readout collapses splice-junction signal into the surrounding window mean and therefore matches the exon-dominant window mean to within rounding, consistent with a tokenisation-induced loss of single-base resolution rather than a model-class disagreement.

**Table A6.**
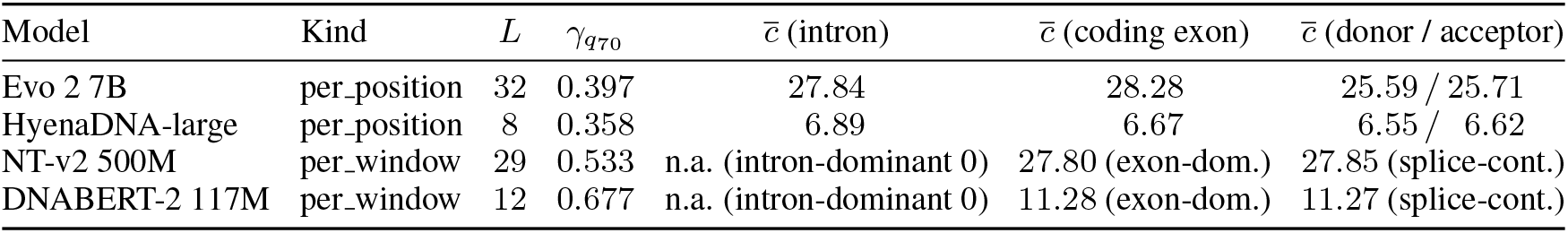
Per-context mean settling depth across the four models on the identical chr22 12,978-window set. Causal-LM models recover donor/acceptor *<* intron at depth-normalised scale; MLM models report per-window estimates with no per-position separation.

#### Pairwise rank concordance

Table A7 shows the pairwise Spearman *ρ* of per-window mean settling depth across the four models. The pattern is two-tier: within-family correlations are positive (*ρ*_Evo2,Hyena_ =+0.516; *ρ*_NT,DNABERT_ =+0.663), whereas every cross-family pair is weakly negative. This is the empirical basis for the “replication within per-bp causal LMs, tokenisation-bound for MLMs” claim of §3.4.

**Table A7.** Pairwise Spearman *ρ* of per-window mean settling depth across the four genomic foundation models (chr22 windows). Within-family correlations are positive; cross-family correlations are weakly negative.

|  | Evo 2 7B | HyenaDNA-large | NT-v2 500M | DNABERT-2 |
| --- | --- | --- | --- | --- |
| Evo 2 7B | 1.000 | +0.516 | −0.119 | −0.188 |
| HyenaDNA-large | +0.516 | 1.000 | −0.287 | −0.166 |
| NT-v2 500M | −0.119 | −0.287 | 1.000 | +0.663 |
| DNABERT-2 | −0.188 | −0.166 | +0.663 | 1.000 |

#### Visual summaries

The per-context bar chart (Fig. A1) shows the donor/acceptor *<* intron inequality directly for the per-bp causal LMs and the absence of separation for the tokenised MLMs. Fig. A2 consolidates the rank-concordance heatmap, the depth-normalised donor-vs-intron comparison, and a schematic of the two-tier structure into a single panel.

**Figure A1.**
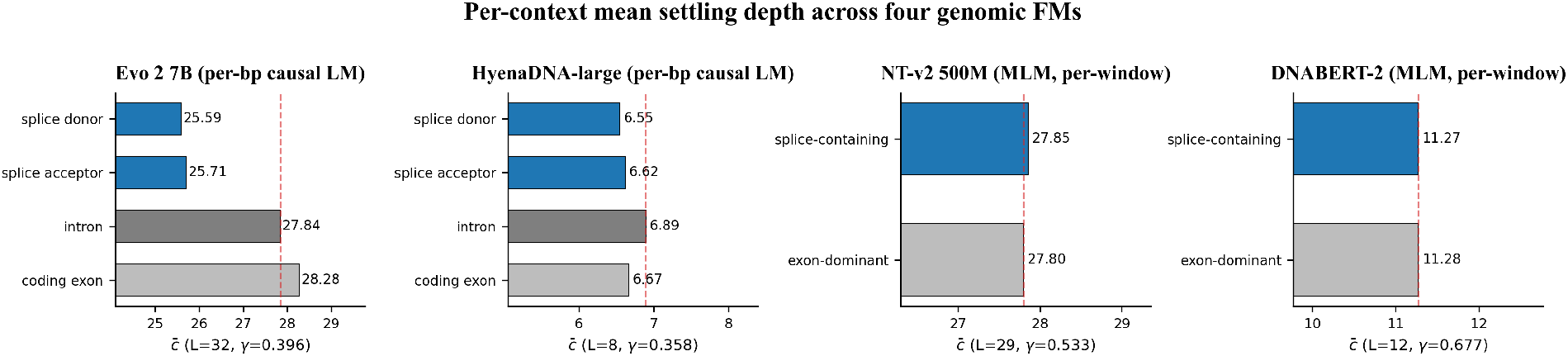
Per-context mean settling depth across the four genomic foundation models. Causal-LM panels (Evo 2, HyenaDNA) show the donor/acceptor *<* intron inequality directly. MLM panels (NT-v2, DNABERT-2) show no separation at tokenised window resolution.

**Figure A2.**
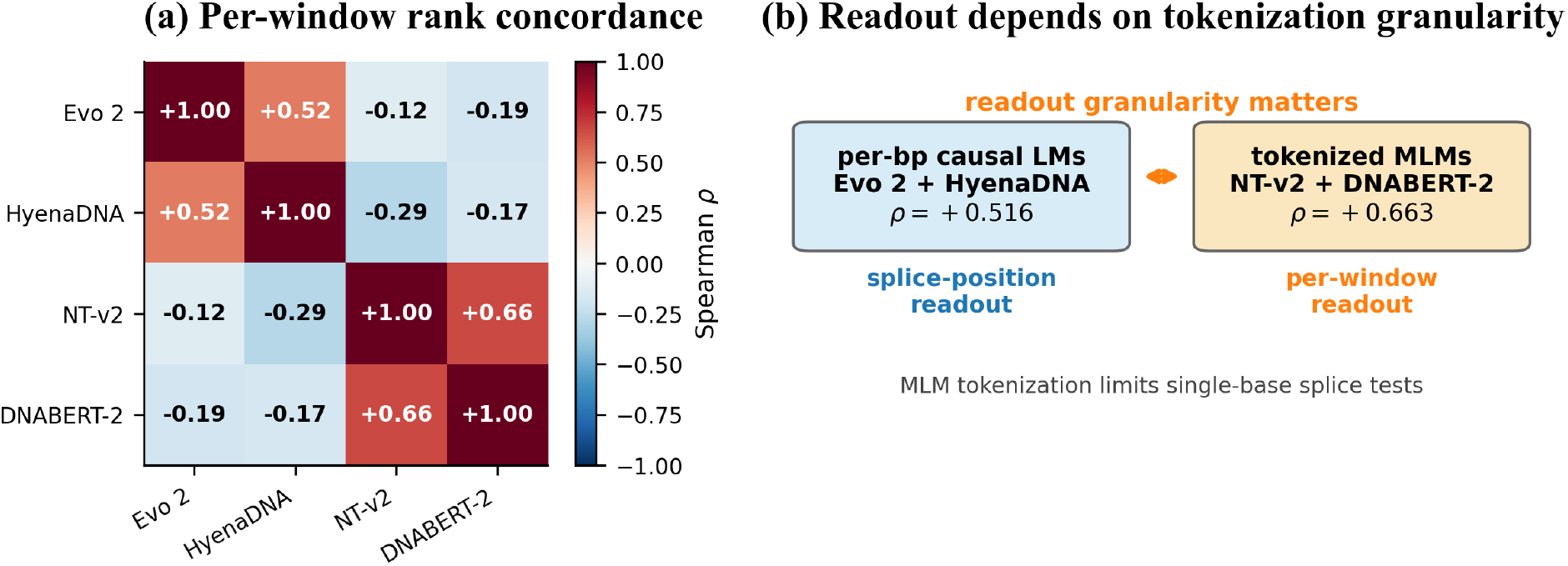
Cross-architecture two-tier structure (numerical detail in Tables A6, A7). **(a)** Pairwise Spearman *ρ* of per-window mean settling depth across four genomic foundation models on chr22, showing the within-family positive block-diagonal vs. cross-family weak-negative off-diagonal structure. **(b)** Schematic of the two-tier structure with the within/cross-family *ρ* values, summarising the “per-bp causal LMs replicate, tokenised MLMs are bound by readout granularity” interpretation.

### C. Variant Pathogenicity: AUROC Summary, Per-Layer Ablation, and Panels

This appendix supports the trajectory-information verification in §3.3. The cohort is the 8,008 ClinVar P/LP-vs-B/LB SNVs across 15 cancer-associated genes; the splitter is 10-fold stratified cross-validation with seed 42, and the LOGO-CV column reports leave-one-gene-out generalisation across the 14 evaluable genes. We treat AUROC strictly as an empirical validation that the 32-d Δ*D*_cos_ trajectory carries information beyond any single tap; we do not position GDTR as a clinical scorer (§3.3).

#### Per-layer ablation

Table A9 reports single-layer logistic-regression AUROC at representative taps for both lenses. The two lenses agree on the U-shape of layer-wise discriminative mass but disagree on which tap is sharpest: JSD concentrates mass at the canonical tap *ℓ* = 29, whereas cosine spreads mass across many taps and accordingly produces a much larger 32-d-vector-vs-best-single-tap gap.

#### Two readings of one lens: why settling and variant scoring favour different taps

Settling depth and variant scoring read the same cosine lens in two different ways, and this is what makes them favour different layers. Settling reads the *absolute* distance *D*_cos_(*ℓ, t*) as a level crossing: it is degenerate at any tap whose representation already matches the reference, so once layer 30 rotates the residual stream into *h*_norm_ the absolute distance collapses to a narrow floor (*D*_cos_(30, *h*_norm_) ≈ 0.31, not 0) and can no longer separate tokens. Variant scoring instead reads the *difference* Δ*D*_cos_(*ℓ, t*) as a sensitivity: subtracting the shared post-rotation floor is precisely what isolates the allele-specific residual, so the signal is sharpest exactly where the absolute distance is least informative. The two readings therefore predict opposite layer-optimality, which is why the single-layer cosine AUROC is 0.604 at *L*^⋆^ = 29 but 0.729 at *ℓ* = 30: the gap reflects one quantity being read before the rotation step and the other after it, not that *L*^⋆^ = 29 is a worse tap. The reference here is *h*_norm_, the post-final-RMSNorm state, not *h*_30_ itself; what saturates at layer 30 is the across-token spread of the absolute distance, which is harmless for the Δ-based sensitivity but fatal for level-based settling. The *ℓ* = 31 AUROC inherits the same 0.729 because layer 31 is a residual passthrough (Tab. A1); it is reported for completeness rather than as new information.

#### ROC, DeLong, and per-gene panels

Fig. A3 provides the four diagnostic panels referenced above. ROC curves (a) confirm the ranking from Table A8; the Paired DeLong tests in (b) show that Δ*D*_cos_ adds significant information beyond Evo 2 ΔLL (*p* = 3.6×10^*−*15^); (c) is the per-layer ablation visualised across all 32 taps, and (d) is the leave-one-gene-out AUROC across the 14 evaluable genes (uniformly ≥ 0.77).

**Figure A3.**
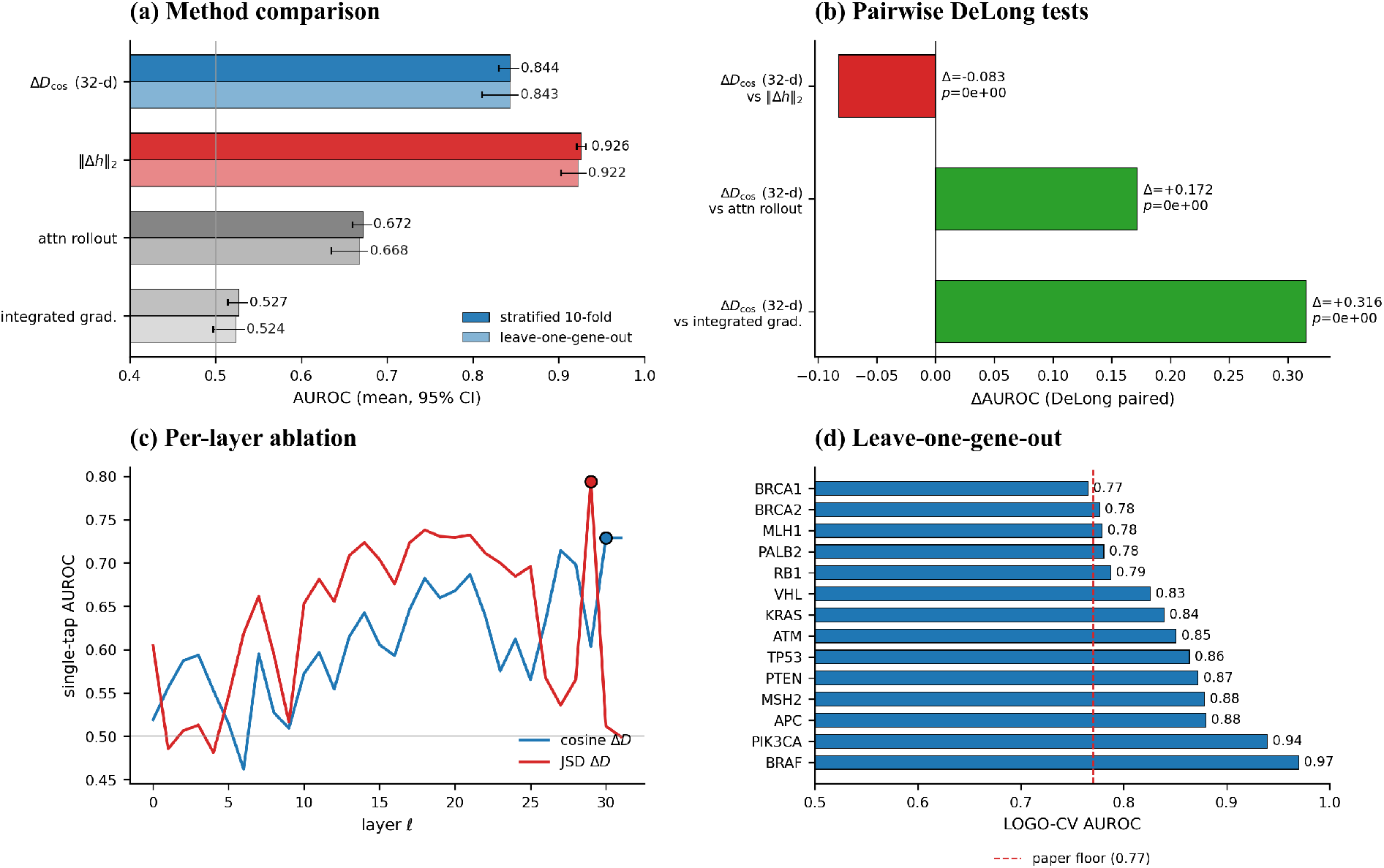
Variant pathogenicity discrimination on 8,008 ClinVar P/LP vs B/LB SNVs across 15 cancer-associated genes (numerical summary in Table A8). **(a)** ROC curves for the four scoring features. **(b)** Paired DeLong tests: Δ*D* adds significant information beyond Evo 2 ΔLL (*p* = 3.6 *×* 10^*−*15^). **(c)** Per-layer single-feature AUROC across all 32 layers – best single tap is *ℓ* = 29 for JSD and *ℓ* = 30 for cosine (the latter is the post-norm rotation layer, not the canonical pre-rotation tap; see App. A.1); the 32-d vector beats either by +0.05 to +0.12. **(d)** Leave-one-gene-out AUROC across 14 evaluable genes is uniformly high (*≥* 0.77).

**Table A8.**
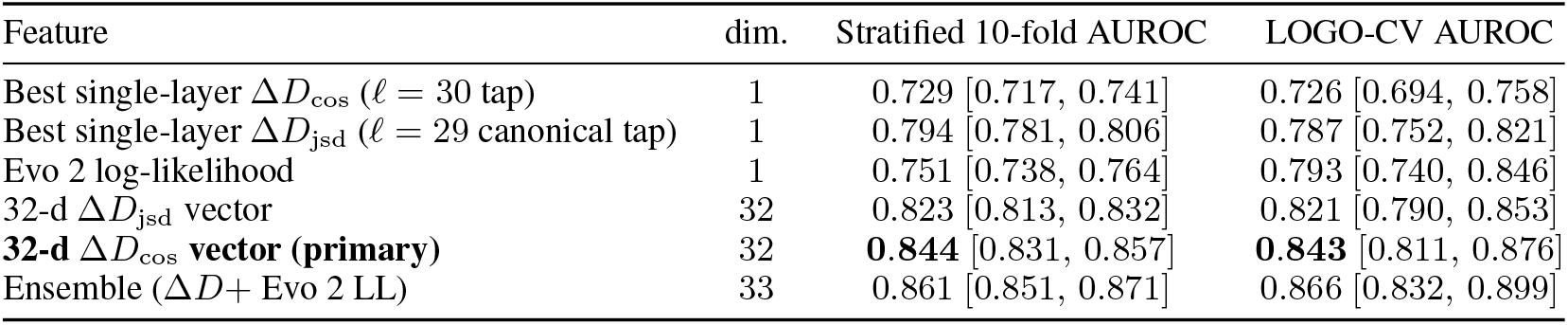
Feature-level AUROC summary. The primary cosine lens reaches 0.844 with the full 32-d trajectory and only 0.729 at its best single tap – a +0.115 gap. Bracketed numbers are 1,000-bootstrap 95% confidence intervals.

**Table A9.** Selected per-layer AUROCs. JSD concentrates discriminative mass at the canonical tap *ℓ* = 29; cosine spreads it across many taps, producing a much larger 32-d-vs-best-single-tap gap.

| Layer | Block type | $\Delta D_{\text{jsd}}$ AUROC | $\Delta D_{\text{cos}}$ AUROC |
| --- | --- | --- | --- |
| 0 | embed | 0.605 | 0.519 |
| 7 | hcs (shallow) | 0.662 | 0.595 |
| 12 | hcm | 0.656 | 0.555 |
| 17 | attn | 0.723 | 0.646 |
| 24 | attn | 0.685 | 0.612 |
| 28 | hcs | 0.565 | 0.698 |
| 29 (canonical tap) | hcm | **0.794** (best JSD) | 0.604 |
| 30 (post-norm-1) | — | 0.512 | **0.729** (best cos) |
| 31 (non-functional, see App. A.1) | — | 0.499 (degen.) | 0.729 (= $h_{30}$) |
| 32-d vector | all | 0.823 | 0.844 |

### D. Splice Anatomy Beyond the Headline Contexts

This appendix supports the cautionary message in §3.2: the splice signal is real, but its internal structure is asymmetric and motif-class-dependent.

#### D.1. Positional Fine-Profile Around Donor / Acceptor

On chr17 the donor profile reaches its mean settling-depth minimum at position +20 bp 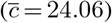 and remains ≥ 1.5 layers below the pooled intronic baseline 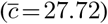 out to ±200 bp. On the same chromosome the acceptor profile reaches its minimum at +50 bp 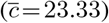 and similarly remains depressed beyond ±200 bp. On chr22 both donor and acceptor minima sit at coordinate 0 (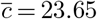 = 23.65 and 23.64 respectively). The asymmetry, with the donor minimum on the exonic side and the acceptor minimum on the intronic side, mirrors the asymmetric splice-grammar features (branch point and polypyrimidine tract) that lie predominantly on the intronic side of acceptors. The full positional profile is shown in Fig. A4; the per-side minima are summarised in Table A10.

**Figure A4.**
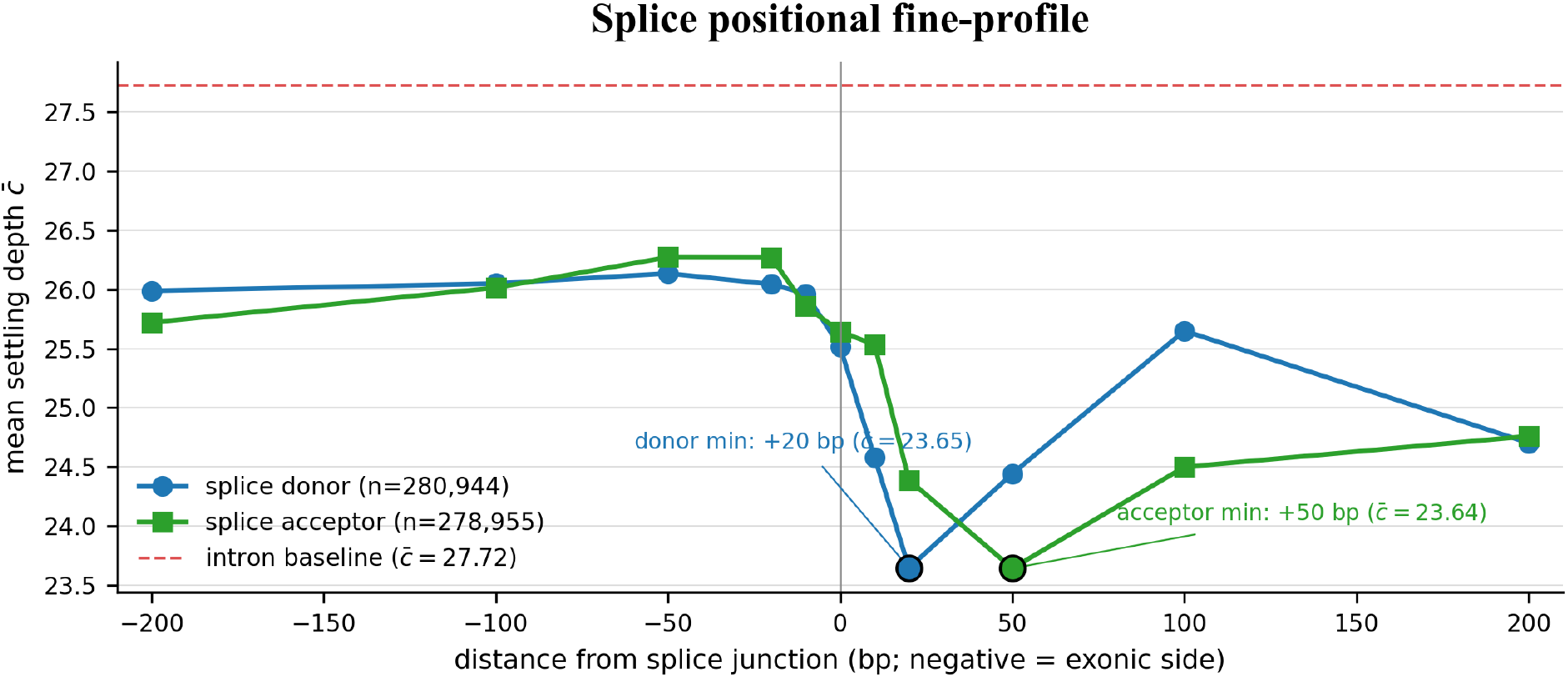
Mean settling depth 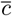 as a function of distance to the nearest splice donor / acceptor, computed on the pooled chr17 + chr22 analysis windows. Both profiles dip well below the intronic baseline (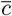 = 27.72, red dashed) within ± 200 bp; donors minimise on the exonic side, acceptors on the intronic side, mirroring the asymmetric splice-grammar context (branch point and polypyrimidine tract on the intronic side of acceptors).

**Table A10.** Per-side minima of the splice positional fine-profile. Pooled chr17 + chr22 analysis windows; positions measured from the splice junction. Donor exonic-side / acceptor intronic-side minima recapitulate the asymmetric splice-grammar context.

| Side | arg-min position | $\bar{c}$ at min | $\bar{c}$ at -100 bp | $\bar{c}$ at +100 bp | $n$ |
| --- | --- | --- | --- | --- | --- |
| splice donor | +20 bp | 23.65 | 26.05 | 25.65 | 280,944 |
| splice acceptor | +50 bp | 23.64 | 26.01 | 24.50 | 278,955 |

#### D.2. Canonical vs. Non-Canonical Splice Motif Breakdown

A finer dissection by motif class, using the genomic dinucleotides immediately downstream of the donor and upstream of the acceptor, reveals an unexpected ordering (Table A11): *non-canonical* splice donors converge *earlier* than the dominant canonical GT-AG donors, while the minor canonical GC-AG class is the deepest of the three. The shallower-than-intron pattern (*c* = 27.72) holds for every splice class, but the within-class ordering is the opposite of a naive “stronger motif implies shallower recognition” prior. The Cohen’s *d* between canonical GT-AG and non-canonical donors is small (≈ 0.05), consistent with the constrained branch-point and polypyrimidine context that flanks non-canonical introns (Wang & Burge, 2008; Burge et al., 1998); we report the magnitudes honestly rather than over-interpret. This ordering is a direct, positive demonstration of arm (b) (context integration) acting independently of motif canonicity. A metric that ranked motifs monotonically by signal strength would have predicted GT-AG *<* GC-AG *<* non-canonical in settling depth. The observed inversion, in which non-canonical donors (e.g. U12-type AT-AC introns) settle earliest because they sit in a tighter, more constrained flank that the model commits to early, is therefore not a contradiction but a clean within-splice instance of the bidirectional reading of §2. Settling depth tracks contextual determinacy, not biological signal strength in any monotone sense. The finding replicates qualitatively on the 8-layer HyenaDNA-large at depth-normalised scale.

**Table A11.** Within-splice motif breakdown by donor dinucleotide. The shallower-than-intron pattern (*c* = 27.72) holds for every splice class, but the within-class ordering is the opposite of a naive “stronger motif implies shallower recognition” prior: non-canonical donors converge earliest, GC-AG latest.

| Donor motif class | $n$ | $\bar{c}$ | vs. canonical GT-AG |
| --- | --- | --- | --- |
| non-canonical (AT-AC, other) | 5,977 | 25.13 | $d \approx -0.05$ (shallower) |
| canonical GT-AG (dominant) | 350,552 | 25.79 | — (reference) |
| canonical GC-AG (minor) | 5,165 | 27.01 | $d \approx +0.10$ (deeper) |
| intron baseline | $\approx 10^7$ | 27.72 | — (deeper than every class) |

#### D.3. Motif and Flank Perturbation Summary

The motif-edit and flank-shuffle controls are reported in the body of the paper at Table 1 alongside the detailed discussion in §3.2. We retain this appendix anchor so that prior references to App. D.3 resolve to the consolidated discussion.

### E. Entropy Decoupling at Genome-Wide Scale

The body of the paper treats *c*(*t*) as decoupled from per-position next-token entropy *H*_t_ at all canonical contexts except 5^*′*^ UTR (§3.1). Here we extend the diagnostic from the 120-window control panel of §3.1 to the full chr22 analysis windows (12,978 windows, 38.9 M analysed positions) to confirm that the decoupling argument is not an artefact of the small control sample. We use the canonical *c*(*t*) from the chr22 forward joined to a per-position *H*_t_ computed from the post-norm logits on the same chr22 sequence. Table A12 reports the per-context joint (*n, c, σ*(*H*_t_), IQR(*H*_t_), *ρ*(*c, H*_t_)).

**Table A12.** Per-context joint distribution of canonical settling depth *c*(*t*) and per-position next-token entropy *H*_t_ on the full chr22 analysis windows (38.9 M positions; central 3 kb of each 12,978 window). *σ*(*H*_t_) and IQR(*H*_t_) measure per-context entropy spread; *ρ*(*c, H*_t_) is Spearman correlation. Contexts are ordered by |*ρ*|.

| Context | $n$ | $\bar{c}$ | $\sigma(H_t)$ | $\text{IQR}(H_t)$ | $\rho(c, H_t)$ |
| --- | --- | --- | --- | --- | --- |
| splice acceptor | 93,883 | 25.69 | 0.496 | 0.944 | -0.224 |
| intron | 20,595,963 | 27.82 | 0.395 | 0.563 | -0.149 |
| splice donor | 94,528 | 25.59 | 0.507 | 0.982 | -0.134 |
| coding exon | 1,643,533 | 28.25 | 0.504 | 1.009 | -0.097 |
| 5' UTR | 74,728 | 29.03 | 0.366 | 0.336 | +0.054 |
| 3' UTR | 610,330 | 27.73 | 0.357 | 0.226 | -0.052 |
| intergenic | 15,821,035 | 28.75 | 0.468 | 0.918 | +0.001 |

Two points are worth noting. First, |*ρ*(*c, H*_t_) | ≤ 0.224 across all seven canonical contexts at this scale; the entropy decoupling argument therefore holds genome-wide. The largest residual couplings appear at splice acceptor (*ρ* = *−*0.224) and intron (*ρ* = *−*0.149); these are negative, indicating that within these contexts higher prediction confidence is associated with shallower settling. This direction is consistent with the splice signal being driven by representational stability rather than by confidence inflation. Second, 5^*′*^ UTR yields *ρ* =+0.054 at *n* = 74,728 positions, smaller than the +0.41 observed on the 120-window control sample. We interpret the control-panel value as a small-sample property of the specific window draw rather than a stable biological feature, and continue to keep 5^*′*^ UTR on a separate axis in Fig. 2 as the most conservative reporting choice. A simple *σ*(*H*_t_)-driven restriction-of-range explanation (whereby narrow entropy spread mechanically suppresses *ρ*) is not supported by the data: 5^*′*^ UTR has the narrowest *σ*(*H*_t_) in the table yet does not produce the smallest |*ρ*|.

### F. Single-Base Promoter Elements

To test whether the splice-site shallow-settling signature of §3.1 generalises to other single-base regulatory anchors, we measure canonical *c*(*t*) at four promoter-element classes on chr22: canonical transcription start sites (TSS, GENCODE v44 Ensembl-canonical), and JASPAR position-weight-matrix hits for TATA-box (MA0108.3, TBP), CAAT-box (MA0060, NFY), and GC-box (MA0079, Sp1). PWM hits are restricted to positions within ± 1.5 kb of a canonical TSS so that each anchor lies in a functional promoter context. Because promoter elements are composition-loaded (GC-box is GC-rich by definition), we report Cohen’s *d* against two baselines: the chr22 intron baseline used elsewhere in the paper, and a GC-content-matched non-promoter chr22 control (5%-wide GC bin, 1:1 matched). Table A13 reports the resulting effect sizes.

**Table A13.** Canonical settling depth at chr22 promoter elements. “vs. intron” is Cohen’s *d* against the chr22 intron baseline; “vs. GC-matched control” is Cohen’s *d* against a chr22 non-promoter control sampled in the same *±*5% GC bin (1:1 matched). For reference, the chr22 splice-donor and splice-acceptor anchors yield *d* = *−*0.367 and *−*0.351 on the same intron baseline.

| Promoter element | $n$ | $\bar{c}$ | $d$ vs. intron | $d$ vs. GC-matched ctrl |
| --- | --- | --- | --- | --- |
| TSS (Ensembl-canonical) | 1,410 | 28.33 | +0.085 | +0.042 |
| TATA-box (MA0108.3, TBP) | 13,560 | 29.14 | +0.219 | +0.027 |
| CAAT-box (MA0060, NFY) | 4,392 | 28.11 | +0.049 | -0.121 |
| GC-box (MA0079, Sp1) | 3,901 | 29.56 | +0.287 | +0.313 |

The splice-shallow signature does not generalise to single-base promoter elements. None of TSS, TATA, CAAT, or GC-box settles shallower than the intronic baseline; TATA and GC-box settle measurably deeper (*d* = +0.22, +0.29 vs. intron). Under the GC-content-matched control, only CAAT-box exhibits a small shallower-settling shift (*d* = −0.12), and GC-box remains robustly deep (*d* = +0.31). TATA and TSS are statistically indistinguishable from their composition-matched controls (|*d*| ≤ 0.05), so the apparent depth shift versus intron in those rows is largely a composition effect rather than a residual-stream signature of motif detection. We read this as direct support for arm (b) of the bidirectional metric: at promoter-element grammars, where Evo 2 must integrate CpG-island and TF-binding-site context, *c*(*t*) shifts deeper rather than shallower. The splice-shallow signal of §3.1 therefore reflects a splice-specific motif-detection arm (arm (a)) rather than a generic short-motif phenomenon.

#### F.1. Per-Context Aggregation Consistency Check

Context-level summaries in the body use per-position *c*(*t*) pooled over windows: 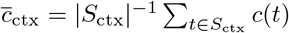. As a consistency check, we recomputed the context-level ranking using per-region mean *c* (one value per annotated region rather than per position). The Spearman correlation between the position-pooled and region-pooled per-context *c* vectors is ≥ 0.97 on both chr22 and chr17, and the rank ordering of the seven contexts is preserved. We report *d* throughout because it normalises by pooled within-context standard deviation and so absorbs within-region variability by construction; both aggregation schemes lead to the same qualitative ordering.

### G. Extended Limitations

This appendix expands the four-kind limitation summary in §4.

#### G.1. Direction-only readout

The cosine lens is scale-invariant, so anything carried in the residual norm ∥*h*_ℓ_(*t*)∥ is invisible to *c*(*t*). Under a superposition view (Elhage et al., 2022; Bricken et al., 2023), the norm reflects how strongly features are active, so two tokens that share a direction but differ in norm get the same settling depth. We use cosine on purpose: Evo 2’s residual norms grow with depth, and a raw-distance lens would confuse directional commitment with that growth. The choice isolates the directional signal and gives up the magnitude. Whether the discarded magnitude matters for settling is testable: at layer 29, the spread of ∥*h*_29_(*t*) ∥ across tokens, and whether it adds anything to *c*(*t*) for variant scoring (App. C), would measure what cosine leaves out. Cosine is also not strictly magnitude-neutral, because a few outlier dimensions can dominate it (Timkey & van Schijndel, 2021); confirming that Evo 2’s stream has no such dimensions, or standardising before the lens, would put *c*(*t*) on firmer ground. This is the same axis as the absolute-versus-differential split in §3.3: the absolute level saturates to a floor, which is fine for the differential variant signal but lossy for the level-based settling readout.

#### G.2. Composition and the off-manifold shuffle

The main standing confound is nucleotide composition. Local base composition (GC content, dinucleotide frequencies) varies systematically with annotation class, so a depth signal that aligns with composition gradients risks being a composition artefact rather than a representational signature. Our entropy decoupling (§3.1, App. A.3 and E) and GC-matched promoter controls (App. F) speak to this. The splice-donor effect survives entropy regression, where *d* strengthens from −0.452 to −0.583 on the residual, and it is splice-specific rather than a generic short-motif effect. The control is incomplete in two places. The §3.2 flank shuffle keeps composition fixed, so a composition-driven model would show a muted shuffle effect rather than none. And App. F shows the confound is real for promoter elements, where TATA’s *d* = +0.22 against intron drops to +0.03 once GC is matched. We read the splice depth signal as composition-robust and treat the promoter and flank-shuffle contrasts as composition-entangled.

The flank-shuffle of §3.2 also has two readings that the binary contrast cannot separate. Under *reduced integration*, depth tracks how much flanking grammar the model integrates, so removing grammar settles *c* earlier. Under *early simplification*, the model treats the shuffled flank as composition-matched noise and commits to the lone GT. Both predict the same direction; they differ only if early simplification is a discontinuous response to noise rather than the smooth limit of graded integration, and a graded-degradation series settles it: shuffle a growing fraction of the flank, or shuffle at growing distance from the junction. A monotone shift in *c* supports reduced integration; a step change supports a separate response to noise. The shuffle is also off-manifold, since dinucleotide-shuffled flanks are out of distribution for a model trained on natural genomes, so it reads out-of-distribution behaviour alongside on-sequence integration. The within-splice ordering in App. D.2 is the on-manifold version of the same test, where non-canonical donors settle earlier through a naturally tighter flank, so we treat it as the primary evidence for context integration and the shuffle as its controlled but out-of-distribution complement.

### H. Data and Code Availability Statement

#### Code and data availability

The integrated analysis pipeline, source code, configuration files, and figure-generation scripts that underlie every numerical result in this paper are publicly available at https://github.com/darejinn/gDTR. All experiments use random seed 42 for cross-validation splits and bootstrap resamples. The Evo 2 model is locked to `arcinstitute/evo2` 7b at HF revision SHA `bda0089f92582d5baabf0f22d9fc85f3588f6b58` (weights MD5 `359ef88ccac2a62644035578de8a7db4)`. Versioned external data sources comprise: GRCh38 primary assembly (UCSC, MD5 locked); GENCODE v44 GTF (per-chromosome filtered and persisted as a gffutils SQLite database); PhyloP 100-way (UCSC); ENCODE SCREEN v3 cCRE catalog (ELS subset); RepeatMasker hg38; GTEx v8 cis-eQTL pairs `(Whole Blood, Brain Cortex, Liver`, and `Lung` tissues, unioned); GWAS Catalog v1.0; and ClinVar archived release dated April 18, 2026. The reference software stack consists of `torch 2.4.1+cu124, evo2 0.3.0, vortex 1.0.8, transformer-engine 2.14.0, transformers 4.49.0, scipy 1.13`, and `scikit-learn 1.4.` Hardware: a single NVIDIA H200 (141 GB). Total compute: approximately 20 GPU-hours end-to-end.

#### Data responsibility and ethics statement

The genomic coordinates and clinical variant datasets analyzed in this study were obtained from publicly available repositories under each repository’s stated license. No restricted human-subject data, identifiable private health information, or individual-level genotype records were acquired or processed. All ClinVar records used here refer to aggregate submitter-classified variant calls without patient identifiers. The integrated pipeline complies with the data-governance frameworks of each upstream repository (UCSC, ENCODE, GTEx, GWAS Catalog, NCBI ClinVar) and uses an archival snapshot (ClinVar release: April 18, 2026) so that all numerical results in this paper remain exactly reproducible from that snapshot regardless of future repository updates.

